# Long-read amplicon denoising

**DOI:** 10.1101/383794

**Authors:** Venkatesh Kumar, Thomas Vollbrecht, Mark Chernyshev, Sanjay Mohan, Brian Hanst, Nicholas Bavafa, Antonia Lorenzo, Robert Ketteringham, Kemal Eren, Michael Golden, Michelli Faria Oliveira, Ben Murrell

**Affiliations:** Department of Medicine, University of California, San Diego, La Jolla, CA, USA; Department of Biology, University of California, San Diego, La Jolla, CA, USA; University of Cape Town, Cape Town, Western Cape, South Africa; Department of Statistics, University of Oxford, Oxford, United Kingdom

**Keywords:** Next generation sequencing, Amplicon sequencing, Amplicon denoising, Long reads

## Abstract

Long-read next generation amplicon sequencing shows promise for studying complete genes or genomes from complex and diverse populations. Current long-read sequencing technologies have challenging error profiles, hindering data processing and incorporation into downstream analyses. Here we consider the problem of how to reconstruct, free of sequencing error, the true sequence variants and their associated frequencies. Called “amplicon denoising”, this problem has been extensively studied for short-read sequencing technologies, but current solutions do not appear to generalize well to long reads with high indel error rates. We introduce two methods: one that runs nearly instantly and is very accurate for medium length reads (here ~2.6kb) and high template coverage, and another, slower method that is more robust when reads are very long or coverage is lower.

On one real dataset with ground truth, and on a number of simulated datasets, we compare our two approaches to each other and to existing algorithms. We outperform all tested methods in accuracy, with competitive run times even for our slower method.

Fast Amplicon Denoising (FAD) and Robust Amplicon Denoising (RAD) are implemented purely in the Julia scientific computing language, and are hereby released along with a complete toolkit of functions that allow long-read amplicon sequence analysis pipelines to be constructed in pure Julia. Further, we make available a webserver to dramatically simplify the processing of long-read PacBio sequences.

## Introduction

The Pacific Biosciences platform allows complex populations of long DNA molecules to be sequenced at reasonable depth. This has been used to study diverse viral populations (1–5) and microbial communities (6, 7), and much more. PacBio SMRT sequencing generates extremely long reads (some >80kb), with very high error rates (~15%) (8). However, this length can be traded for accuracy. By ligating hairpin adapters that circularize linear DNA molecules, the sequencing polymerase can make multiple noisy passes around single molecules, and these can be collapsed into Circular Consensus Sequences (CCS) that have much higher accuracy (9).

When sequencing amplicons of a fixed length, the number of passes (ie. the total raw read length divided by the amplicon length) is a primary determinant of the accuracy of a CCS. The raw read length distribution has a long right tail, which means that the number of passes around each molecule, and consequently the CCS error rates, can vary substantially. Here we confine our discussion to these CCS reads.

A critical feature of PacBio sequences is a high homopolymer indel rate. (3) show that, for a 2.6kb amplicon, with a post-filtering error rate of 0.5%, 80% are indel and 20% substitution errors, and the indel errors are concentrated in homopolymer regions, increasing in rate with the length of the homopolymer. While high indel rates can be computationally challenging to deal with, since sequence alignment can be slow, they are favorable from a statistical perspective, because the errors appear in predictable places, making them more correctable (10).

Amplicon denoising (11–17) refers to a process that takes a large set of reads, corrupted by sequencing errors, and attempts to distill the noiseless variants and their frequencies. This has been extensively studied for short-read sequencing technology, but these approaches do not always generalize well to longer reads.

It is helpful to distinguish between two sequencing regimes: short and accurate (SA), and long and inaccurate (LI), and PacBio sequencing datasets can span both of these. For a given error rate, the probability of an observed read being noise free decreases exponentially with read length, and the error rate determines how precipitous this decline is (see Figure 1). For short, accurate reads, we can expect to have many noiseless representative reads in our dataset. Indeed, many Illumina amplicon denoising strategies (11, 18) rely on this, and amount to simply identifying these reads using their relative abundance information. Shorter PacBio reads fall into this category as well. However, as the amplicon length increases, not only are there more opportunities for error, but the number of passes around each molecule decreases, increasing the per base error rate. There may be variants that simply do not have any noiseless representatives, forcing us to abandon these “read-selection” strategies of amplicon denoising in this long, inaccurate regime. We can only hope to reconstruct the noiseless reads, by identifying a set of noisy reads that originate from the same variant, and averaging out their noise.

**Fig. 1.**
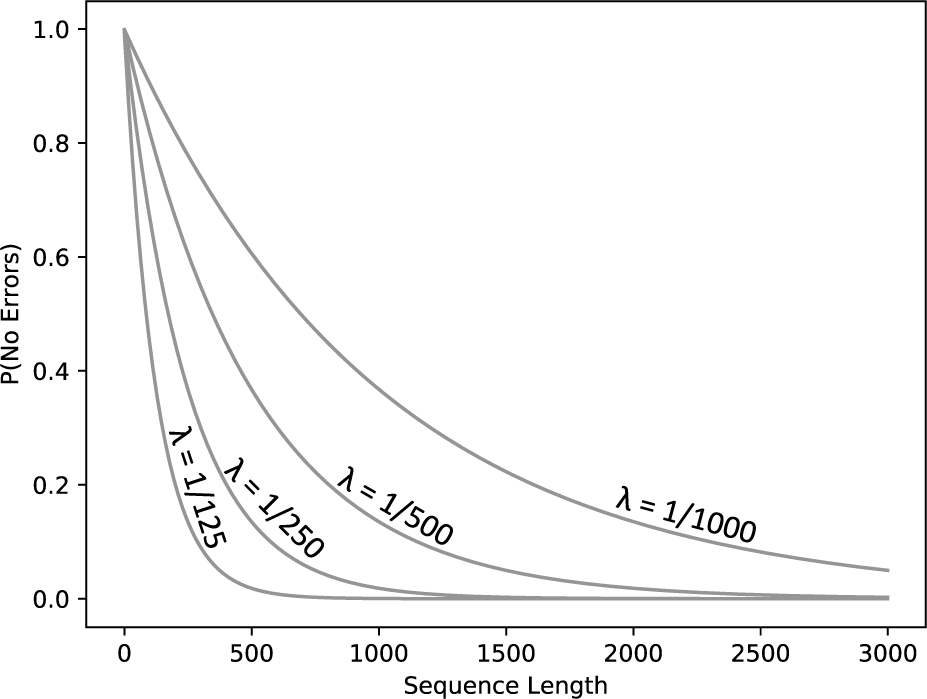
Under simple assumptions, the probability that a sequence will have no errors decreases exponentially with the sequence length, and the slope of this decrease is determined by the per-base error rate.

Existing approaches to this have used off-the-shelf clustering tools to render approximate reconstructions of the underlying population (3, 19, 20), but these are not built for purpose and can, as we show here, be improved upon substantially.

Our strategy mirrors this distinction, with one tool (FAD) that operates in the SA regime, and one (RAD) that operates in the LI regime. Both are implemented entirely in Julia, an emerging language for scientific computing.

## Approach

We present two methods: the Fast Amplicon Denoiser (FAD), and the Robust Amplicon Denoiser (RAD). FAD is designed for cases where an appreciable number of sequences are expected to be error free, and these can reliably serve as our inferred templates, avoiding any form of clustering or consensus calls, and exploiting abundance and neighborhood information to keep or reject templates. This method performs better for shorter amplicons, higher quality sequencing, and better read-per-template coverage.

RAD is more complex, and designed for cases where very few reads are error free. This can occur in PacBio amplicon sequence when either amplicons are very long, with fewer passes per molecule, or for short movie lengths, reducing raw read lengths, or for older sequencing chemistries. RAD works in stages. First, we employ a kmer-domain clustering approach, inspired by a non-parametric Bayesian procedure (21, 22) to partition reads into clusters, followed by a recursive cluster refinement procedure (also in kmer domain).

## Methods

### A. Kmer representation

For both RAD and FAD, we heavily exploit a kmer-based distance calculation. We first convert all sequences to their kmer counts. For all analyses here, *k* = 5 or *k* = 6, representing each sequence as a vector of integers of length 4^*k*^. We then seek to approximate the pairwise edit distance between two sequences using these kmer frequency vectors.

While there exist sophisticated distance metrics based on kmer similarity (23), we opt for a simple approach that scales linearly with substitutions for low-divergences. Consider two identical sequences, with identical associated kmer vectors. When a random substitution is introduced, there will typically be ~2k differences between the kmer vectors. So our kmer approximation of edit distance is simply:

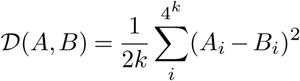

See figure 2 for a demonstration of how this behaves, compared to edit distance. We can optionally scale this distance by dividing by the sequence length, to yield a per-base percentage difference.

**Fig. 2.**
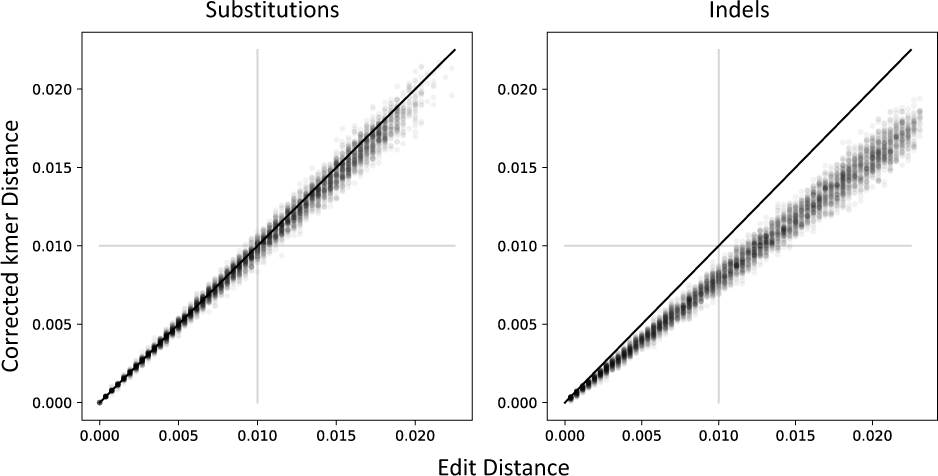
This distance approximates edit distance as mutations are introduced, starting from the 2599bp NL4-3 HIV-1 env sequence. When only substitutions are introduced, edit distance is extremely well approximated. When indels are introduced, our kmer distance underestimates edit distance. This is desirable behavior when the sequencing error process is dominated by indels, because they will be down-weighted in our distance function.

### B. Fast Amplicon Denoising (FAD)

FAD is the simpler of the two algorithms, intended to work in low-noise scenarios. FAD proceeds by de-replicating reads, and sorting them by abundance, ignoring all reads that do not occur at least twice. FAD iterates through each read from largest to smallest, maintaining a set of accepted templates. When the current read is distant from all reads already included in the set by ≥ 1bp (as calculated by our corrected kmer distance), then it is added to the set. If it is within 1bp, then the abundances of the higher frequency template are considered when deciding to keep or discard the lower frequency template. We first, however, correct the abundances by the expected proportion of error-free sequences. We convert the QV scores into error probabilities, and obtain an expected number of errors per sequence. We then evaluate the probability of each sequence having zero errors, and take the mean of this. For our LP135 dataset, this comes to 38%.

We take the most abundant template ≤ 1bp from the current template, and we calculate the p-value for the size of a spurious offspring that differs at one base, under a Poisson error assumption, Bonferroni corrected for the average number of sites in the template. If this is ≤ *α* (default: *α* = 0.01), then we reject the null hypothesis that we would obtain an offspring template this large by chance, and we include this template in the set.

Finally, we take all reads, and assign each read to an accepted template based on the minimum distance under the kmer approximation (using *k* = 5 here, since this method places an emphasis on speed). These are used to compute the final frequencies for all reads.

### C. Robust Amplicon Denoising (RAD)

RAD is intended for high-noise scenarios, where we do not expect sufficient numbers of reads to be noise free for the strategy employed by FAD to succeed. We nevertheless aim to keep the computation time as low as possible, exploiting kmer distances extensively.

#### C.1. Dirichlet process means clustering

We wish to cluster our kmer frequency vectors. We do not know how many clusters we have in advance (ruling out traditional options like k-means), and we need the algorithm to scale well with the input dataset size and the number of clusters. PacBio error rates per read are, however, highly predictable from the quality scores, so we have information about the distance by which a noisy read can differ, by error alone, from its noise-free platonic ideal. Fortunately, the “Dirichlet process means” (DP-means) clustering approach (21) is ideal here.

It is frequently observed that k-means clustering can be derived as an expectation maximization algorithm for a finite mixture of isotropic Gaussians, where the variance of the Gaussians is sent to zero, forcing hard-assignments of elements to clusters (24). Similarly, DP-means can be derived as the limit of sampling procedure for a non-parametric Bayesian Dirichlet process infinite mixture of Gaussians model, where the variance is similarly driven to zero. This yields a surprisingly simple deterministic algorithm that uses a “radius” parameter *λ* to control the number of clusters (21). Briefly, the DP-means algorithm works by maintaining an array of centroids, and passing through the elements one at a time, computing the distance to all cluster centroids: if the distance between the element and any centroid is *< λ*, then assign the element to the cluster with the nearest centroid, and if not, seed a new cluster, using that element as the cluster centroid. After each pass through the elements, recompute the cluster centroids by averaging all the elements that are assigned to them. This iterates until convergence. See (21) for a technical description.

We use this algorithm to cluster our kmer vectors, using the scaled kmer distance, and a radius *λ* = 0.01, which is the error rate we typically use to retain .fastq reads in our data filtering steps.

#### C.2. The triangle inequality

The number of clusters is typically be much lower than the number of reads. After the first DP-means clustering pass yields a set of centroids, we cluster these centroids to construct a set of “meta-centroids”, and we compute, just once, the pairwise distance between all reads and all meta-centroids. Upon each subsequent iteration, we compute the pairwise distance between all current centroids and all meta-centroids, and we use the triangle inequality to avoid computing the read-to-centroid distances when we can deduce that they are *> λ*, reducing computation by a factor that depends on the template diversity.

#### C.3. Fine cluster splitting

Clustering reads using a radius equal to the error filtering cutoff (we often use 1%) can fail to distinguish variants that are very closely related. We therefore introduce a second layer of cluster refinement that directly seeks to split clusters that are different at any bases. Again, for computational efficiency, we remain in the kmer frequency vector domain to avoid sequence alignment.

Consider a cluster of a few closely related variants, each with multiple reads corrupted by sequencing noise (which has errors scattered at random bases). We attempt to suppress the noise by identifying the kmers that differ the most, and cluster just on these, with a very low clustering radius. To avoid splitting on homopolyer errors, we choose M (default M=20) kmers with the largest variance, and search this set of high-variance kmers for kmer pairs that differ by a single homopolymer length edit, discarding these. We take the highest variance remaining N (default N=6) kmers, and run DP-means clustering on this very low dimensional representation of the reads, with euclidean distance, and a default radius of 1. Ie. if any reads differ at more than one of these kmers, we separate them. Please note that 1bp difference should cause at least 6 kmers to differ, so this can split reads that differ by a single base.

This clustering step produces a “candidate” cluster split, which we then decide to accept or reject, using the abundance information of these sub-clusters. If the original cluster gets fragmented into too many small clusters that fall below a size threshold, we reject the split. For this, we use the same Bonferroni corrected Poisson p-value approach as used in FAD. After splitting, we recurse, and continue splitting each sub-cluster until there is no evidence of heterogeneity.

#### C.4. Kmer-seeded alignment consensus

Unlike FAD, the clusters identified by RAD are not expected to have noise-free sequences associated with them. We thus rely on a consensus approach to infer these templates. We start by finding the sequence whose kmer vector is nearest to the cluster average kmer vector, which we take as a draft consensus. We then align all reads, pairwise, to this draft consensus. Using these pairwise alignment coordinates, we run a sliding window over the draft consensus, and when any blocks of this draft reference do not match the most common sub-sequence of the aligned reads, we replace that block of the draft reference with this modal value. We exploit kmer seeding (k=30), and this approximate pairwise alignment algorithm scales linearly with sequence length.

If the amplicon spans a coding sequence, then Rifraf.jl (10) can be used to infer a frame-shift corrected template sequence, as long as a reference sequence with a trusted reading frame is available.

### D. A metric for comparing inferred to true templates

A population can be represented by a set of sequences, and their associated frequencies. We seek a metric that can be used to evaluate the reconstruction accuracy of an algorithm. A useful distance metric for evaluating reconstruction accuracy must be zero when the reconstructed sequences and frequencies are identical to the ground truth, should grow as the divergence between the two sets grows, and should have a meaningful numerical interpretation. One attractive option here is the Earth Mover’s Distance, operating on the matrix of pairwise distances and frequencies. We have previously advocated this (19), but here we expand on this a little. We now refer to this as “Sequence Mutation Distance” (*S M D*), and release SMD.jl, which calculates this metric. A related approach, UniFrac (25, 26) is commonly used to compare microbial communities, but UniFrac computes distances over a phylogeny, where as SMD operates directly on the pairwise distance matrix.

Consider the ground truth sequences *A*, and the inferred templates *B* and a distance matrix *D* wherein *D*_*i,j*_ is the distance between *A*_*i*_ and *B*_*j*_ (here we use edit distance). Construct a flow matrix *F*, which is of the same shape as *D* but *F*_*i,j*_ represents how much of *A*_*i*_ maps onto *B*_*j*_.

*SMD* can be defined as:

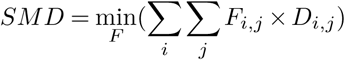

with respect to constraints:

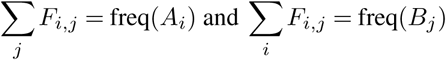

Where *freq*(*X*) is the frequency associated with variant *X*. This SMD score corresponds to the weighted average number of nucleotide changes per sequence required to convert *A* to *B*, finding the (possibly non-unique) minimum by optimizing over *F*. In our implementation, we use the Julia package JuMP.jl (27) to perform this optimization.

This can be interpreted as the total error in the reconstruction per sequence. Indeed, if we compute the *S M D* between noisy sequences and the templates from which they were derived, we obtain a very precise estimate of the empirical error rate, biased only very slightly towards underestimation (see figure 3).

**Fig. 3.**
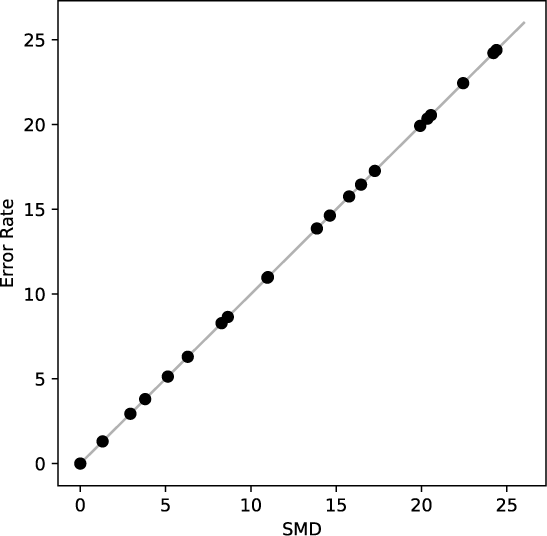
SMD accurately approximates the average error rate, when computed between a set of templates, and a set of sequences that are derived from the templates by some noisy process.

The SMD score is optimized while respecting frequency constraints on both ground truth sequences *A*, and the inferred templates *B*. We can additionally derive two scores of interest: By relaxing the constraint on frequencies of *A*, we get *S M D*_*FP*_, which increases with the extent and frequency of false positives (ie. reconstructed sequences that are absent from the ground truth). Similarly, by eliminating the constraint on *B*, we get *S M D*_*FN*_, a measure of false-negatives, which increases when our reconstructions are missing sequences that are present in the ground truth.

### E. Comparison methods

1. baseline: This refers to the original set of reads with no denoising.
2. VSEARCH: VSEARCH’s cluster-fast is run with an identity threshold of 0.99 (equivalent to our radius threshold of 0.01), and the consensus output is evaluated.
3. USEARCH: USEARCH’s cluster-fast is run with identical parameters to VSEARCH, with an id threshold of 0.99. The “consensus output” is used as inferred templates.
4. deep USEARCH: Similar parameters as the USEARCH method, but with the “max-accepts” parameter set to 300 instead of the default, and the “maxrejects” parameter set to 600 instead of the default, to cause USEARCH to search more aggressively for better matches during clustering.
5. UNOISE: Fastx_uniques is run with a size output to dereplicate reads, followed by UNOISE3, using the “amplicon output” (without chimera filtering) as inferred templates. Please note that the UNOISE documentation asserts that UNOISE is not designed for PacBio data.

All methods were run single-threaded on an AMD Ryzen 7 1700 processor @ 3.0 Ghz.

## Results

We assembled four datasets to compare methods; one real, and three simulated using the PacBio sequence simulator developed in (19):

1. LP135: We used a number of HIV envelope clones available in our lab, all from the same time point, and isolated from the same donor. To construct a ground truth clustering for these reads, we amplified 96 wells using paired forward and backwards primers that uniquely identify the well, sequenced on the RS-II using P6/C4 chemistry, with 6 hour movie lengths. CCS reads were inferred with PacBio’s CCS algorithm (v3.0). From this .fastq sequencing dataset, we first filter at the 1% accuracy threshold (3), partly to guarantee accurate barcode sequencing, and we recover 80 pure clones. The consensus of reads from each well is taken as the ground truth sequence. When inferring templates using RAD, FAD, and other methods, we first trim off the barcodes from the .fastq reads, to ensure the true clustering is obscured. The full dataset had ~18k reads, but we also subsampled datasets of 10k, 5k, and 2k to investigate lower template coverage (which could occur when multiplexing samples).
2. P018 early. This is a simulated dataset. We obtained templates and frequencies from a run of the Full Length Envelope Analysis pipeline (FLEA) (3). The “early” dataset is simulated from the P018 “V06” time point, approximately 6 months post-infection, representing a challenging dataset of low diversity. We include an unfiltered, and a 1% filtered dataset.
3. P018 late. As for “early”, but using the 33 months postinfection templates and frequencies, which had higher diversity. The error profiles for the P018 datasets were generated to match P5/C3 chemistry (the previous generation), and have a higher mean error rate than our real P6/C4 dataset, which impacts the relative performance of the methods. We include unfiltered 1% filtered datasets.
4. 9kb. We simulated low-diversity evolution of 9kb templates, starting from the full-length nl43 plasmid, with random mutations (including indels), generating 32 closely-related templates. Frequencies were simulated from a uniform distribution. We matched the error rates in the simulated reads to those from a 9kb plasmid sequence (data not shown), and these are substantially lower than the ~2.6kb amplicons, primarily due to the amplicon length. Here we include unfiltered, 1% filtered, and 2% filtered datasets.

Template sequence phylogenies and summary statistics for these datasets are depicted in figure 4.

**Fig. 4.**
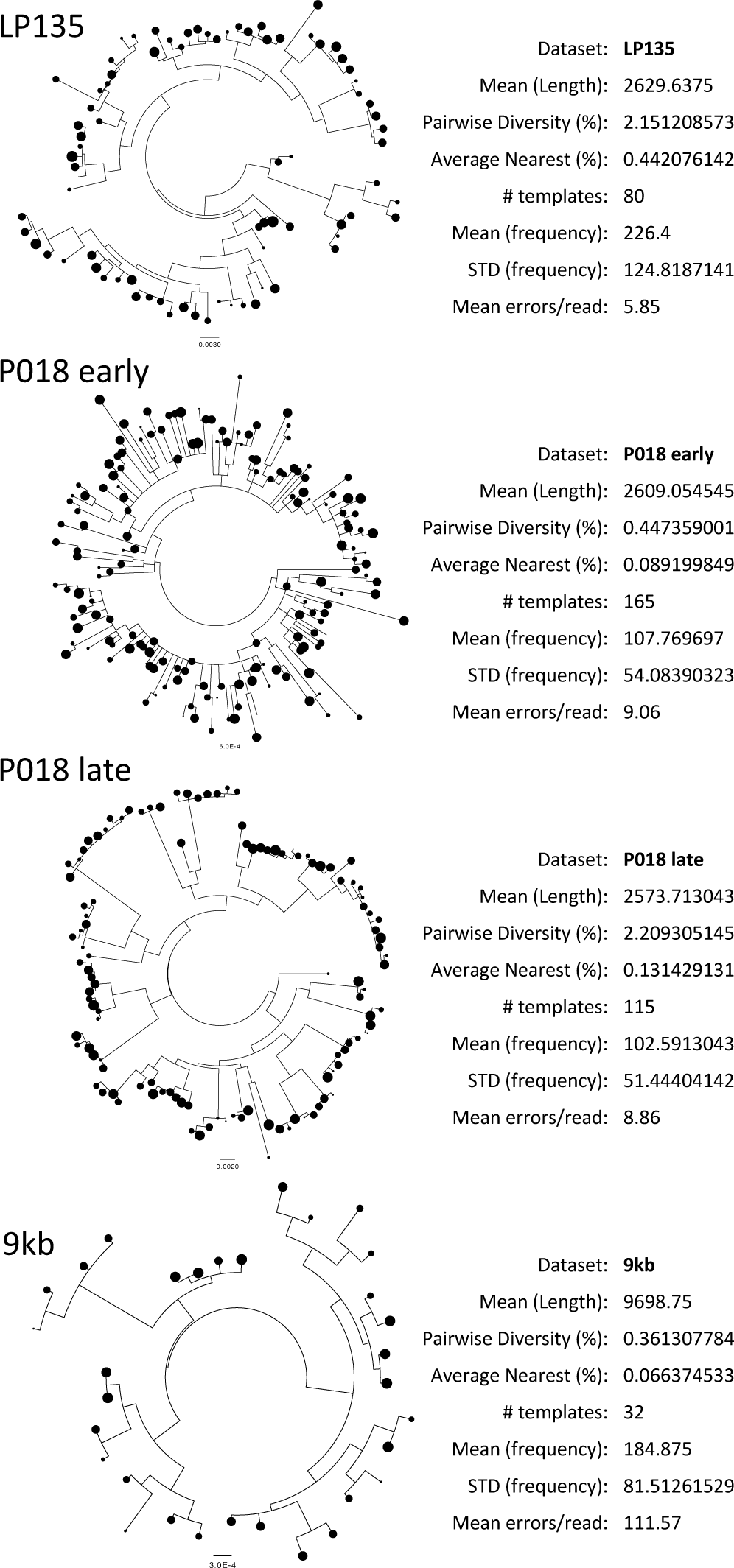
Four test datasets. See main text for more detailed descriptions. Here we show maximum likelihood phylogenies of the template sequences, along with summary statistics: the mean read length, the mean pairwise diversity, the average distance from each template to its closest neighbour, the number of templates, the average template frequency, the standard deviation over template frequencies, and the mean number of errors per read.

### F. Performance

See figure 5 for accuracy (SMD scores) and timing results. These SMD scores are not normalized by sequence length, and can be interpreted as the per-sequence error rate. So an SMD of 1.0 means that there is, on average, 1bp incorrect in each sequence. False positive and false negative SMDs are shown in figures S1 and S2.

**Fig. 5.**
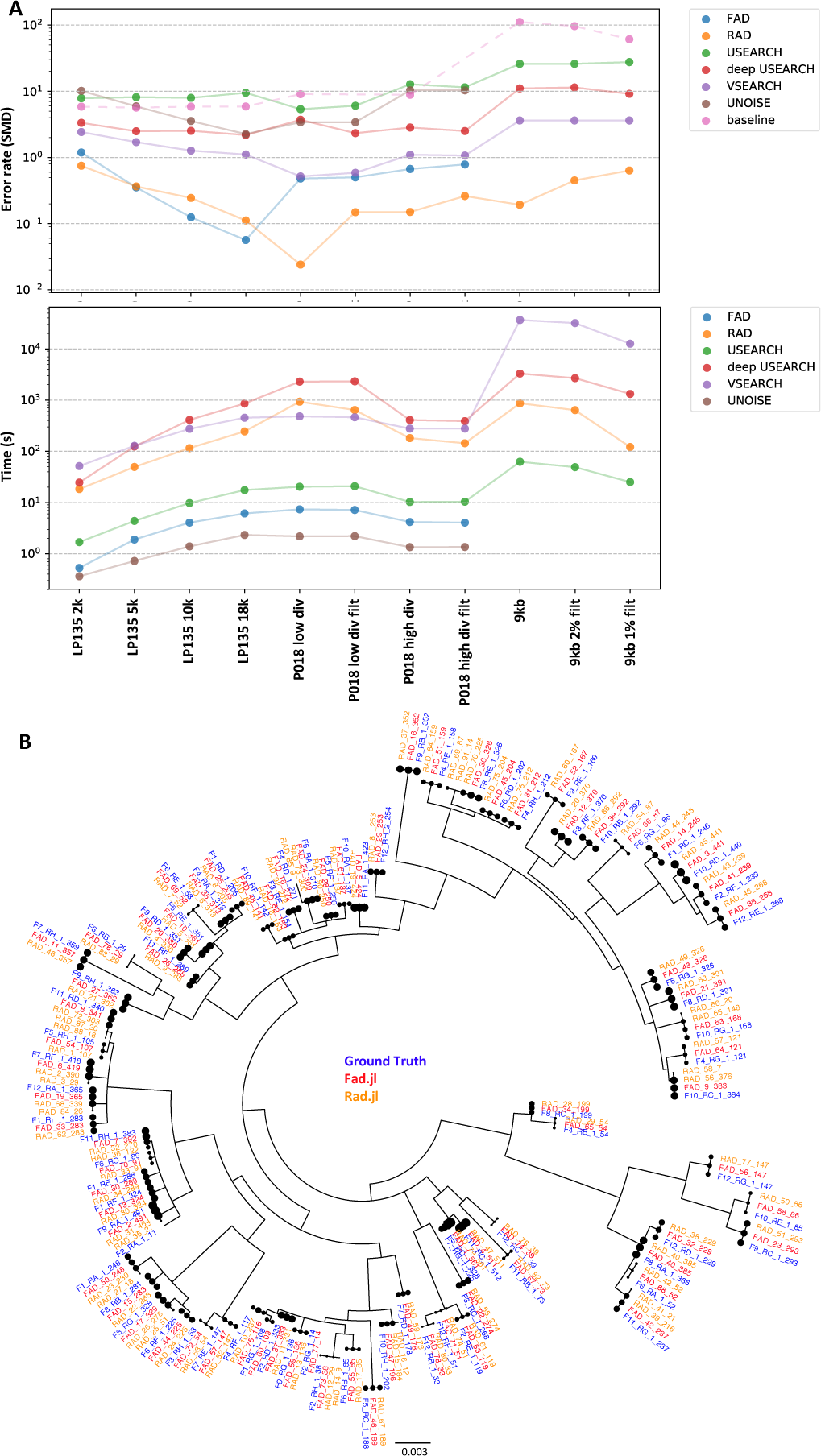
Panel **A**: Error rates (measured by Sequence Mutation Distance) of reconstructions against ground truth for a number of datasets. LP135 is a real sequencing dataset, using primer barcodes to obtain the ground truth clustering. P018 (~2.6kb) comprises reads simulated *in silico* from a set of templates obtained from an HIV+ donor, from a low diversity, early time point, and a later, more diverse, time point. The 9kb dataset comprises a set of closely related templates, with long reads simulated from these, using a higher error rate profile. The dashed “baseline” depicts the SMD scores of the uncorrected reads against the ground truth templates. Also shown are run times of the various methods. Panel **B**: From the LP135 full dataset, we show a phylogeny depicting the ground truth templates, as well as the inferred templates for FAD and RAD.

USEARCH (with default parameters) and UNOISE are fast, but inaccurate. In many cases, USEARCH has an accuracy similarly to the SMD of uncorrected reads (“baseline” in the figure). UNOISE, as expected, is not well suited to these long-read datasets. Deep USEARCH, modified for a more extensive search during clustering, is slower and more accurate than USEARCH. The timing difference can be dramatic: from a minute for USEARCH, to nearly an hour for deep USEARCH, when inferring templates from the unfiltered 9kb dataset. VSEARCH has intermediate accuracy, with SMDs as low as 0.51 for the P018 low-diversity dataset. VSEARCH runs in time comparable to deep USEARCH for 2.6kb datasets, but becomes very slow for 9kb datasets, taking 10 hours on the slowest dataset.

FAD (like UNOISE), does not complete on the 9kb datasets, because it requires an appreciable proportion of error-free reads. FAD is extremely fast on all 2.6kb datasets, never taking longer than 10 seconds. FAD is also the most accurate method on the full LP135 dataset, with an SMD of 0.057 (which translates to a per-base error rate of 1 in ~46000). As expected, FAD’s performance degrades as the template coverage decreases, and as the error rates increase (the simulated 2.6kb datasets had higher error rates than LP135). RAD is always faster than deep USEARCH, and faster than VSEARCH in all-but-two datasets, and has especially well controlled run times for the 9kb datasets. RAD, however, stands out as being consistently accurate across all datasets, with results close to FAD in the low-noise LP135 datasets, but with clearly superior results across the noisier regimes. The closest competitor, VSEARCH, has substantially higher SMD scores than RAD on all datasets, with accuracies ranging from 3.2x to 18.7x worse.

Please note that these results should not be taken as a criticism of USEARCH or VSEARCH’s clustering, or UNOISE, as these algorithms were not designed with this problem in mind.

## Conclusion

We have presented two algorithms, FAD and RAD, for denoising long PacBio amplicons. While we intend to use these tools primarily for applications in virology, which has motivated our choice in datasets, there is no reason why they cannot be used in any long-read amplicon sequencing domain, especially for metagenomics (eg. 16S).

With this paper we release four Julia packages: NextGenSeqUtils.jl, Rad.jl, DPMeansClustering.jl, and SMD.jl, which should be a helpful contribution to the Julia next generation sequencing ecosystem. We also provide a web server (https://tools.murrell.group/denoise) for convenient analyses. A .fastq CCS file is uploaded, and filtering options can be selected. RAD is run, and the inferred templates, as well as a number of visualizations (see figure 6), are provided.

**Fig. 6.**
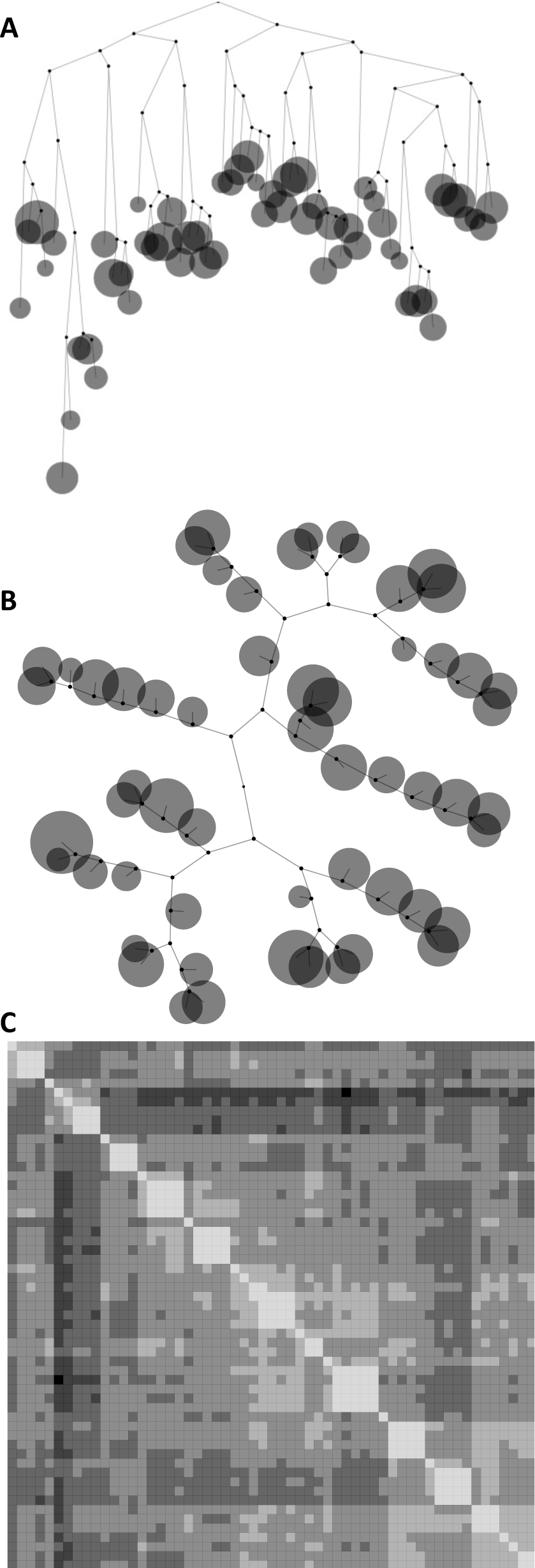
Interactive visualization of inferred templates and their frequencies is available in multiple layouts on the RAD/FAD webserver. Neighbour joining (28) phylogenies are inferred from fast corrected kmer distance matrices, and displayed in traditional phylogeny format (A), as well as D3 force directed graph layout (B). Sequence names are shown interactively. We also display distance matrices (C). Together, these allow a rapid assessment of the diversity and population structure of the inferred template sequences.

The algorithms could potentially be improved along multiple dimensions:

1. Automatically determining the optimal method for amplicon denoising: we currently use a simple heuristic to choose which of RAD or FAD should be used. This uses the QV scores to obtain an estimate of the expected number of error free reads, and it uses the proportion of identical reads. If both of these are sufficiently high, use FAD, but if either is low, we recommend RAD. The details of what counts as high or low require further exploration on additional datasets.
2. Test other kinds of sequencing data: future work should compare amplicon denoising methods on datasets from a wider range of sources, spanning a range of length and template diversity.
3. Using error rates when clustering: since the error rates are highly predictable, the distance between a read and a centroid could be adjusted by the expected error in each, which could result in more accurate clustering.
4. Using error rates when splitting: per-base error rates could also be exploited during cluster splitting for both FAD and RAD, potentially improving accuracy.
5. Parallelization: We could gain run-time improvements by parallelizing some components of our model. The simplest of these would be the RAD consensus step, where each consensus can be executed on a different thread.

Additional extensions may be domain specific. For example, chimera filtering (29–31) is not useful in domains like HIV, where extensive biological recombination produces the same signals as artificial chimeras. However, this could be useful in other domains, and should be implemented.

PacBio have released a “Long Amplicon Analysis” (LAA) tool. This tool runs directly on the raw sequence data, however. This prevents comparison on any of our simulated datasets, where we simulate the .fastq reads directly, and even our real PacBio dataset had primer barcodes trimmed at the CCS level, preventing comparison. Future work should find a way to compare LAA to these approaches where ground truth is available.

The advent of accurate long-read denoising approaches shifts the developmental burden away from data processing. Going forward, the primary impediment to extending the length of amplicons that can be sequenced is the design of PCR strategies that can successfully amplify very long templates.

## ACKNOWLEDGEMENTS

Library preparation for SMRT sequencing was performed by the Translational Virology Core at the UC San Diego Center for AIDS Research, with special thanks to Caroline Ignacio and Gemma Caballero. SMRT sequencing was conducted at the IGM Genomics Center, University of California, San Diego, La Jolla, CA. Thanks to Sasha Murrell for copy editing.

This work was supported by the National Institute Of Allergy And Infectious Diseases of the National Institutes of Health under Award Numbers R00AI120851, R01AI120009, and UM1AI068618, the National Institute on Drug Abuse under Award Number R21DA041007, and in part by the University of California, San Diego Center for AIDS Research (P30AI036214). K.E. was supported in part by R21AI115701. The content is solely the responsibility of the authors and does not necessarily represent the official views of the National Institutes of Health. MFO was supported by Conselho Nacional de Desenvolvimento Científico Tecnológico (CNPq)-Brazil.

---

**Table S1.** Single-threaded timing results and SMD scores for all tested datasets.

| Dataset | Method | Time | SMD | SMD_FP | SMD_FN |
| --- | --- | --- | --- | --- | --- |
| LP135_2000 | FAD | 0.529428005 | 1.19 | 0.0 | 0.7805 |
| LP135_2000 | RAD | 18.34718585 | 0.750451427 | 0.015523285 | 0.442 |
| LP135_2000 | UNOISE | 0.363973141 | 10.15344068 | 0.0 | 6.325 |
| LP135_2000 | USEARCH | 1.691693068 | 7.839674207 | 1.025658339 | 4.6495 |
| LP135_2000 | deep USEARCH | 24.61717916 | 3.339864407 | 0.848587571 | 2.076 |
| LP135_2000 | VSEARCH | 51.52917194 | 2.424851197 | 0.350057013 | 1.0615 |
| LP135_5000 | FAD | 1.898756027 | 0.353 | 0.0 | 0.2822 |
| LP135_5000 | RAD | 49.55263019 | 0.365213846 | 0.016806723 | 0.1516 |
| LP135_5000 | UNOISE | 0.724266052 | 5.923901299 | 0.0 | 1.5628 |
| LP135_5000 | USEARCH | 4.379507065 | 8.148839892 | 0.860107817 | 2.7488 |
| LP135_5000 | deep USEARCH | 124.194135 | 2.490797184 | 0.760008799 | 1.489 |
| LP135_5000 | VSEARCH | 128.1675749 | 1.7074 | 0.186666667 | 0.822 |
| LP135_10000 | FAD | 4.062191963 | 0.1248 | 0.0069 | 0.0635 |
| LP135_10000 | RAD | 115.500721 | 0.244969081 | 0.02721633 | 0.1496 |
| LP135_10000 | UNOISE | 1.397036076 | 3.555361582 | 0.0 | 0.6545 |
| LP135_10000 | USEARCH | 9.790716171 | 7.949061126 | 0.8086386 | 3.2107 |
| LP135_10000 | deep USEARCH | 410.9284868 | 2.529141943 | 0.71510222 | 1.3111 |
| LP135_10000 | VSEARCH | 274.4071872 | 1.2691428 | 0.161719096 | 0.443 |
| LP135_18112 | FAD | 6.140394926 | 0.056647527 | 0.006404594 | 0.010103799 |
| LP135_18112 | RAD | 244.904155 | 0.111601287 | 0.03318975 | 0.027385159 |
| LP135_18112 | UNOISE | 2.334758997 | 2.258735458 | 0.0 | 0.114675353 |
| LP135_18112 | USEARCH | 17.54733491 | 9.48204451 | 0.508407756 | 1.450640459 |
| LP135_18112 | deep USEARCH | 857.4190199 | 2.183995122 | 0.544159372 | 1.105675795 |
| LP135_18112 | VSEARCH | 453.11621 | 1.108383671 | 0.080107134 | 0.239068021 |
| P018_low_D | FAD | 7.381646156 | 0.480317175 | 0.0 | 0.429310539 |
| P018_low_D | RAD | 930.7800851 | 0.024111227 | 0.0 | 0.014059161 |
| P018_low_D | UNOISE | 2.189254999 | 3.412146332 | 0.0 | 1.597064447 |
| P018_low_D | USEARCH | 20.39851213 | 5.36600672 | 0.050422902 | 3.902766843 |
| P018_low_D | deep USEARCH | 2298.212697 | 3.707657171 | 0.820386618 | 2.070408278 |
| P018_low_D | VSEARCH | 482.140717 | 0.51757209 | 0.035695961 | 0.15465077 |
| P018_low_D_filt | FAD | 7.172053814 | 0.501284493 | 0.0 | 0.429310539 |
| P018_low_D_filt | RAD | 640.839519 | 0.149100384 | 0.0 | 0.097233157 |
| P018_low_D_filt | UNOISE | 2.214979172 | 3.412146332 | 0.0 | 1.597064447 |
| P018_low_D_filt | USEARCH | 20.86764693 | 6.044407546 | 0.965805406 | 4.393993926 |
| P018_low_D_filt | deep USEARCH | 2328.007018 | 2.329603861 | 0.240870527 | 0.905522438 |
| P018_low_D_filt | VSEARCH | 462.745703 | 0.586916761 | 0.04795373 | 0.128669441 |
| P018_high_D | FAD | 4.176362991 | 0.670113579 | 0.0 | 0.633073402 |
| P018_high_D | RAD | 181.33832 | 0.150246184 | 0.002798507 | 0.120867944 |
| P018_high_D | UNOISE | 1.350667 | 10.34358931 | 0.0 | 6.253432785 |
| P018_high_D | USEARCH | 10.27629089 | 12.77710231 | 0.133790738 | 1.7104594 |
| P018_high_D | deep USEARCH | 408.661937 | 2.834875035 | 0.328613629 | 1.064248178 |
| P018_high_D | VSEARCH | 277.8375401 | 1.099081516 | 0.020323427 | 0.239447364 |
| P018_high_D_filt | FAD | 4.062068939 | 0.78546184 | 0.0 | 0.633073402 |
| P018_high_D_filt | RAD | 143.951633 | 0.261548057 | 0.001048584 | 0.106373962 |
| P018_high_D_filt | UNOISE | 1.359381914 | 10.34358931 | 0.0 | 6.253432785 |
| P018_high_D_filt | USEARCH | 10.40896988 | 11.48160262 | 0.062843296 | 1.48008137 |
| P018_high_D_filt | deep USEARCH | 388.9146309 | 2.509755897 | 0.248885202 | 1.04602475 |
| P018_high_D_filt | VSEARCH | 277.8611 | 1.070193933 | 0.011554393 | 0.23673504 |
| 9kb | FAD | 5.053105116 | NaN | 0.0 | NaN |
| 9kb | RAD | 867.3528638 | 0.193204868 | 0.0 | 0.189317106 |
| 9kb | UNOISE | 2.392202139 | NaN | 0.0 | NaN |
| 9kb | USEARCH | 62.53130293 | 26.0050563 | 9.884057971 | 23.83536173 |
| 9kb | deep USEARCH | 3294.691269 | 11.0451932 | 5.82962963 | 5.594996619 |
| 9kb | VSEARCH | 36730.41921 | 3.623855472 | 0.211838006 | 0.640804598 |
| 9kb_filt2 | FAD | 4.358752966 | NaN | 0.0 | NaN |
| 9kb_filt2 | RAD | 637.2588739 | 0.450119459 | 0.0 | 0.189317106 |
| 9kb_filt2 | UNOISE | 2.134759903 | NaN | 0.0 | NaN |
| 9kb_filt2 | USEARCH | 49.09791207 | 25.99045984 | 7.318681319 | 23.76200135 |
| 9kb_filt2 | deep USEARCH | 2682.655685 | 11.42585245 | 5.905109489 | 5.950811359 |
| 9kb_filt2 | VSEARCH | 31920.33632 | 3.623855472 | 0.211838006 | 0.640804598 |
| 9kb_filt1 | FAD | 1.947170019 | NaN | 0.0 | NaN |
| 9kb_filt1 | RAD | 121.6790071 | 0.636113083 | 0.0 | 0.054428668 |
| 9kb_filt1 | UNOISE | 1.025439024 | NaN | 0.0 | NaN |
| 9kb_filt1 | USEARCH | 25.10640097 | 27.6137593 | 10.33333333 | 25.46771467 |
| 9kb_filt1 | deep USEARCH | 1322.334428 | 9.144750493 | 5.266313933 | 5.835192698 |
| 9kb_filt1 | VSEARCH | 12609.67987 | 3.62229784 | 0.211838006 | 0.640804598 |

**Fig. S1.**
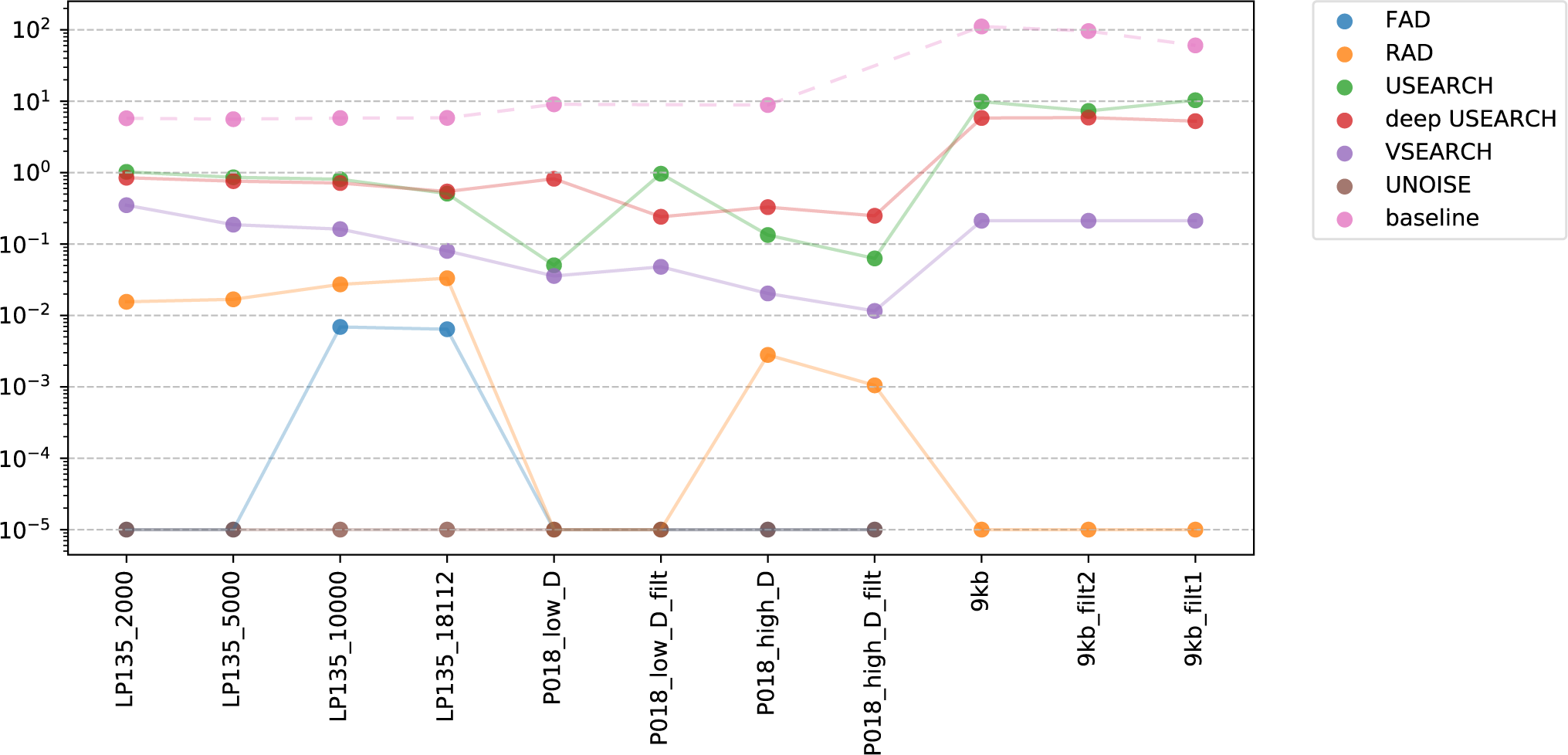
False positive error rates (measured by *S M D*_*FP*_) of reconstructions against ground truth for a number of datasets. Values of 0 are set to 10^*−*5^ for the log transform.

**Fig. S2.**
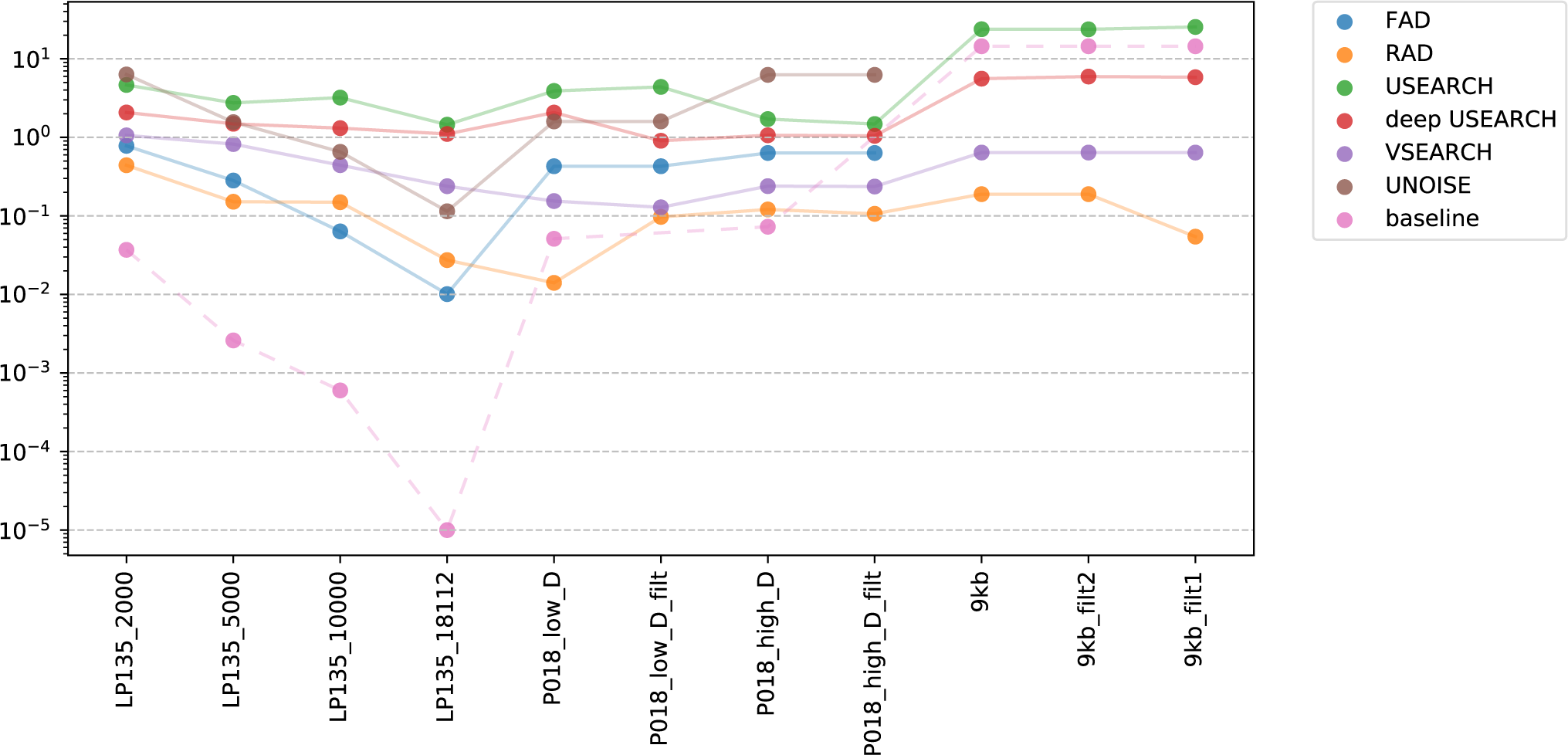
False negative error rates (measured by *S M D*_*F N*_) of reconstructions against ground truth for a number of datasets. Values of 0 are set to 10^*−*5^ for the log transform.

